# Ageing reshapes the resting-state network landscape of the human brain

**DOI:** 10.64898/2026.09.07.749858

**Authors:** N. Kudriavtsev, M. Rosso, G. Fernández-Rubio, E. Serra, M.H. Andersen, M. L. Kringelbach, P. Vuust, L. Bonetti

**Affiliations:** Center for Music in the Brain, Department of Clinical Medicine, Aarhus University & The Royal Academy of Music, Aarhus/Aalborg, Denmark; Department of Cognitive Neuroscience, Faculty of Psychology and Neuroscience, Maastricht University, Netherlands; Centre for Eudaimonia and Human Flourishing, Linacre College, University of Oxford, Oxford, United Kingdom; Department of Psychiatry, University of Oxford, Oxford, United Kingdom; Danish Research Centre for Magnetic Resonance, Department of Radiology and Nuclear Medicine, Copenhagen University Hospital - Amager and Hvidovre, Hvidovre, Denmark; Faculty of Health and Medical Sciences, University of Copenhagen, Copenhagen, Denmark

**Keywords:** Ageing, Resting-state, Brain networks, Alpha slowing, MEG, Generalised eigendecomposition (GED)

## Abstract

Ageing alters brain rhythms and large-scale functional organisation, but whether these changes reflect isolated effects or a coordinated reconfiguration remains unclear. We applied Frequency-resolved Network Estimation via Source Separation (FREQ-NESS) to source-reconstructed resting-state MEG from 164 healthy younger and older adults across two independent datasets. FREQ-NESS revealed that ageing reshaped the frequency-resolved organisation of whole-brain networks. Older adults showed a slower alpha-network peak and a flatter decay of network prominence from alpha into beta frequencies, together with changes in entropy across the spectrum. These spectral changes were accompanied by shifts in the spatial expression of dominant networks along the lateral, posterior-anterior, and inferior-superior spatial axes. Rather than following a uniform anatomical shift, the direction of age differences varied across frequencies. Together, these findings show that healthy ageing is associated with a reshaped configuration of endogenous brain networks, characterised not simply by a uniform loss of network organisation, but by selective reconfiguration of the prominence and spatiotemporal expression of whole-brain activity across frequencies.

## Introduction

The global population is ageing at an accelerating rate. By the late 2070s, the number of people aged 65 years or older is projected to exceed the number of children under 18 worldwide (United Nations Department of Economic and Social Affairs [UN DESA], 2024). Understanding how the brain changes across later stages of life is therefore an increasingly important scientific and societal challenge. Yet healthy ageing is not a uniform process. Individuals differ widely in cognitive trajectory, structural integrity, and in the capacity to tolerate age-related neural change (Cabeza et al., 2018; Turrini et al., 2023). Furthermore, ageing bears an impact on various levels of the brain, affecting cellular homeostasis, synaptic plasticity, neuromodulation, vascular and metabolic function, and the integrity of axons and myelin (Bishop et al., 2010; Turrini et al., 2023). These changes ultimately constrain the timing and coordination of activity across large neural populations, raising the question of how local and distributed alterations combine to reshape whole-brain communication. Classical electrophysiological signatures, such as slowing of the individual alpha rhythm, may therefore represent isolated consequences of ageing or different manifestations of a broader reorganisation of endogenous brain activity across frequencies and space (Scally et al., 2018; Stacey et al., 2021; Hinault et al., 2023; Merkin et al., 2023).

Resting-state electroencephalography (EEG) and magnetoencephalography (MEG) recordings have identified several recurrent signatures of healthy ageing, including changes in oscillatory power, dominant frequency, aperiodic spectral structure, connectivity, and signal complexity (Babiloni et al., 2016; Cassani et al., 2018; Ishii et al., 2017; Fernández-Rubio et al., 2025). Among these, slowing of the dominant alpha rhythm is one of the most consistent findings. Older adults typically show a lower individual alpha peak frequency, but this effect is distinct from a reduction in alpha power (Merkin et al., 2023; Scally et al., 2018). This distinction is important because within conventional frequency band analyses, it is commonly assumed that the same spectral boundaries apply across individuals and age groups. Scally et al. (2018) found lower upper-alpha power and connectivity in older adults when conventional bands were used, but no group differences were observed when activity was evaluated at single participant’s individual alpha peak. Spectral estimates are further affected by the broadband aperiodic background, often referred to as 1/f spectral component, which reflects the non-oscillatory decrease in signal power with increasing frequency and can obscure or bias estimates of narrowband oscillatory peaks. Separating periodic and aperiodic components attenuated the age difference in alpha peak power reported by Merkin et al. (2023), while the slowing of alpha peak frequency remained significant. Ageing has also been associated with a flatter aperiodic exponent (Voytek et al., 2015), with age-related differences in 1/f spectral activity also reported in more recent EEG work (Criscuolo et al., 2025). However, estimates of aperiodic parameters depend on the fitted frequency range, treatment of oscillatory peaks, and other modelling choices (Donoghue et al., 2020, 2022; Merkin et al., 2023). Together, these findings show that preprocessing decisions can conflate changes in peak position, oscillatory amplitude, and background spectral structure, therefore challenging the classical established effects of ageing and urging for a more holistic approach.

A complementary literature has examined ageing at the level of large-scale brain networks. Resting-state fMRI commonly shows weaker connectivity within functional systems, stronger coupling between networks, and lower segregation or modularity in older adults (Chan et al., 2014; Deery et al., 2023; Geerligs et al., 2015). These changes are often described as increased integration, but this interpretation is metric-dependent. Greater between-network coupling can coexist with lower local or global efficiency and weaker hub organisation (Deery et al., 2023). Electrophysiology adds a further dimension because network organisation can be resolved by frequency and at millisecond timescales. However, functional connectivity and network architecture findings are not always consistent across modalities, partly because of differences in methodological approaches (Maestú et al., 2019). Recent MEG studies analysing source-reconstructed data show that age effects vary across cortical regions, frequency ranges, and transient network states rather than following a single monotonic pattern (Andersen et al., 2026; Bonetti et al., 2024; Gohil et al., 2026; Malvaso et al., 2026; Quinn et al., 2025; Ranasinghe et al., 2025; Ruuskanen et al., 2026). Multimodal work further suggests that age-related functional reorganisation is shaped by structural connectivity, but in a region- and frequency-specific manner: multilayer analyses combining diffusion MRI and MEG have shown that structure-function coupling in alpha-band temporal and parietal networks is associated with cognitive performance in older adults (Jauny et al., 2024). The emerging view is therefore one of selective reorganisation under biological constraint, rather than uniform loss across all neural measures.

Ageing has also been associated with changes in the spatial distribution of functional brain activity. One influential example is the posterior-anterior shift in ageing, which describes reduced posterior and relatively greater anterior task-related activation in older adults (Davis et al., 2008). However, this framework was developed primarily from task-evoked fMRI, and an anterior shift is not inherently compensatory. Instead, compensation requires evidence that altered recruitment responds to neural demands and supports preserved performance (Cabeza et al., 2018). More broadly, age-related reorganisation may be expressed along several anatomical dimensions rather than a single posterior-anterior axis. A frequency-resolved source-space approach can therefore test whether the dominant spatial configuration of endogenous activity shifts along lateral, posterior-anterior, or inferior-superior gradients, without assigning an adaptive or maladaptive function to those changes.

These limitations have encouraged attempts to describe electrophysiological ageing through higher-level properties of neural dynamics rather than isolated spectral markers. One such direction has been the use of entropy-based measures, which can capture aspects of signal variability and nonlinear organisation that may remain undetected by conventional power-spectrum or connectivity analyses (Cacciotti et al., 2024; Keshmiri S., 2020). However, the resulting measures are heterogeneous: sample entropy, multiscale entropy, permutation entropy, Katz fractal dimensionality, Higuchi fractal dimensionality, and Lempel-Ziv complexity quantify different features of temporal regularity, scale dependence, waveform geometry, or sequence compressibility, and depend strongly on scale, frequency, data length, and parameter selection (Lempel & Ziv, 1976; Katz, 1988; Higuchi, 1988; Richman & Moorman, 2000; Costa et al., 2002; Bandt & Pompe, 2002). A more explicitly multivariate approach was recently proposed by Diambra et al. (2026), who combined principal component analysis with entropy of the normalised covariance eigenvalues to quantify global synchronisation across multiple EEG channels, thereby moving beyond pairwise measures of neural interaction. This approach illustrates how the distribution of variance across concurrent components can provide information that is not available from individual channels or connections alone. Given that oscillatory changes are among the most prominent but method-dependent signatures of ageing, a data-driven framework that jointly resolves the frequency-specific prominence, multivariate component structure, and spatial organisation of whole-brain activity may provide a more integrated account.

## Materials and methods

The goal of the study was to characterise how healthy ageing reshapes the frequency-resolved dynamics and spatial organisation of whole-brain networks reconstructed from resting-state MEG. To do so, we analysed source-reconstructed resting-state MEG data from two independent datasets of healthy younger and older adults **(Figure 1a-d)**. We used Frequency-resolved Network Estimation via Source Separation (FREQ-NESS) to estimate, for each participant and frequency, the prominence and spatial configuration of dominant brain networks **(Figure 1e-i)**. We then asked whether ageing reshaped this network landscape at several complementary levels: (1) the dominant resting-state network landscape; (2) the individual alpha-network peak; (3) the alpha-to-beta decay of the leading network; (4) the entropy of the network landscape; and (5) the spatial gradients of frequency-resolved networks.

**Figure 1.**
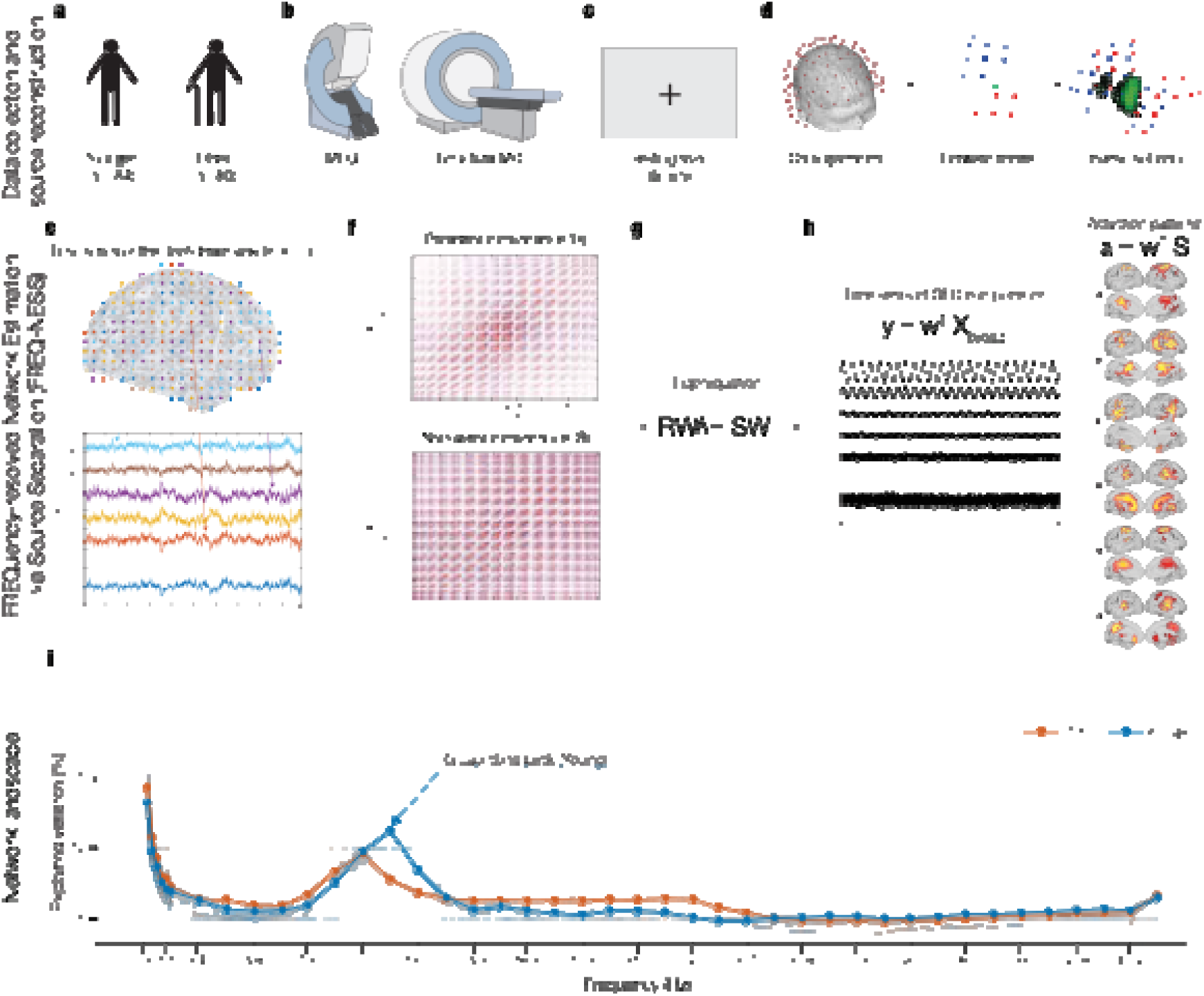
FREQ-NESS analysis of healthy ageing. **(a)** Open eyes resting-state MEG data were analysed from 164 healthy participants, comprising 84 younger and 80 older adults drawn from two independent datasets. **(b)** MEG recordings and structural T1-weighted MRI scans were acquired for anatomical source reconstruction. **(c)** For each participant, an approximately five-minute (304 seconds) resting-state MEG segment was selected for analysis. **(d)** Structural MRI and MEG data were co-registered, after which forward and inverse models were computed to reconstruct source-level neural activity. **(e)** Beamforming yielded a source-level data matrix, containing the time series of 3559 brain voxels. **(f)** At each frequency, a broadband reference covariance matrix, (R), was computed from the unfiltered source-level data, and a narrowband covariance matrix, (S), was computed after frequency-specific filtering. **(g)** Generalized eigendecomposition (GED) was used to identify spatial filters that maximised narrowband covariance relative to broadband covariance. The resulting eigenvectors defined the spatial filters, whereas the corresponding eigenvalues quantified the variance explained by each frequency-specific component. (h) Applying the spatial filters to the source-level data yielded one time series for each GED component; the corresponding spatial activation patterns were obtained by multiplying the filters by the narrowband covariance matrix. **(i)** The normalised eigenvalue of the leading GED component was evaluated across frequencies to define each participant’s frequency-resolved component landscape. Faint lines represent individual participant landscapes, and the superimposed lines and points represent the group-level profiles for younger and older adults. The individual alpha-network peak was defined as the frequency at which the leading component explained the greatest proportion of variance within the predefined alpha-range search window. The lower-frequency portion of the component landscape is displayed for illustration. This leading-component landscape provided the basis for subsequent analyses of the individual alpha-network peak and alpha-to-beta decay, while the full GED eigenspectrum and corresponding activation patterns were additionally used to quantify component entropy and spatial gradients, respectively.

### Participants and study design

We analysed resting-state MEG data from 164 participants drawn from two independent datasets of healthy younger and older adults **(Figure 1a)**. Dataset 1 included 77 participants, comprising 37 younger adults (mean age = 21.89 years, SD = 2.05) and 40 older adults (mean age = 67.50 years, SD = 5.46). Dataset 2 included 87 participants, comprising 47 younger adults (mean age = 22.70 years, SD = 2.48) and 40 older adults (mean age = 69.54 years, SD = 6.65). The combined sample, therefore, included 84 younger and 80 older adults. Younger adults were defined as participants younger than 27 years, whereas older adults were defined as participants older than 55 years.

Participants were included if the recordings passed the preprocessing and source-reconstruction procedures described below. Recordings were excluded in cases of technical issues during acquisition, or incomplete preprocessing. The two datasets were first analysed separately and were then combined for the main analyses, allowing us to assess whether the principal ageing effects were consistent across independent samples.

Dataset 1 was approved by the Institutional Review Board at Aarhus University [Case No. DNC-IRB-2021-012]. Dataset 2 was approved by the Ethics Committee of the Central Denmark Region [De Videnskabsetiske Komitéer for Region Midtjylland, Ref. 1-10-72-127-23]. All procedures were carried out in accordance with the Declaration of Helsinki. All participants gave informed consent before starting the experimental procedure.

### MEG and MRI data acquisition

MEG recordings were acquired in a magnetically shielded room at Aarhus University Hospital, Aarhus, Denmark, using an Elekta Neuromag TRIUX system with 306 channels, consisting of 204 planar gradiometers and 102 magnetometers **(Figure 1b)**. Recordings were sampled at 1000 Hz with online analogue filtering between 0.1 and 330 Hz. Before each MEG session, the participant’s head shape and the positions of four head-position indicator coils were digitised relative to three anatomical landmarks using a Polhemus Fastrak 3D digitizer. The head-position indicator coils were used to continuously track head position during acquisition and to support movement correction during preprocessing. Bipolar electrodes were used to record electrooculographic (EOG) and electrocardiographic (ECG) activity, which were later used to identify eye-blink and heartbeat-related artefacts.

Structural MRI scans were collected using a CE-approved 3T Siemens MRI scanner at Aarhus University Hospital. T1-weighted anatomical images were acquired using an MPRAGE sequence with fat saturation at a spatial resolution of 1.0 x 1.0 x 1.0 mm. Sequence parameters were: echo time = 2.61 ms, repetition time = 2300 ms, reconstructed matrix size = 256 x 256, echo spacing = 7.6 ms, and bandwidth = 290 Hz/Px. MEG and MRI recordings were conducted on separate days. Only resting-state MEG recordings were analysed in the present study.

### MEG preprocessing

MEG processing followed a standard preprocessing and source-reconstruction pipeline used in previous studies from our group (Bonetti et al., 2022; Fernández-Rubio et al., 2022; Rosso et al., 2025; Malvaso et al., 2026). Raw MEG data were first processed with MaxFilter to attenuate external noise sources and correct for head movement. Signal-space separation was applied using the spatiotemporal extension, together with movement compensation based on the continuous head-position information. Data were downsampled by a factor of four, from 1000 Hz to 250 Hz, and a correlation limit of 0.98 was used between the inner and outer subspaces. The resulting sampling rate allowed us to analyse frequencies up to 97.6 Hz, remaining below the Nyquist frequency of 125 Hz, while improving the computational feasibility of whole-brain source-level covariance estimation.

The data were then converted to Statistical Parametric Mapping format and further processed in MATLAB using custom-built code together with the OHBA Software Library, FieldTrip, FSL, and SPM (Friston et al., 2007; Oostenveld et al., 2011; Smith et al., 2004; Quinn et al., 2022). Continuous MEG recordings were visually inspected to identify gross artefacts. Independent component analysis, as implemented in OSL, was then used to remove eye-blink and heartbeat artefacts. The original MEG signal was decomposed into independent components, and each component was correlated with activity recorded from the EOG and ECG channels. Components showing markedly stronger correlations with the EOG or ECG than the remaining components were labelled as candidate artefacts. These candidate components were then visually inspected in terms of their activation time courses and scalp topographies to confirm the expected stereotyped patterns generated by eye blinks or heartbeats. Components were removed only when both the correlation-based criterion and visual inspection indicated an ocular or cardiac origin. The signal was then reconstructed by back-projecting the remaining components into MEG sensor space.

For each participant, the preprocessed continuous resting-state recording was segmented into a single approximately five-minute epoch **(Figure 1c)**. To maintain consistency with the original FREQ-NESS implementation and the analysis code used here, the analysed segment corresponded to 304 s of data sampled at 250 Hz.

### Anatomical source reconstruction

To provide a physiologically interpretable characterisation of whole-brain network activity, we reconstructed the anatomical sources underlying the MEG signal. Source reconstruction was performed using beamforming, combining custom-built code with routines from OSL, SPM, FieldTrip, and FSL (Friston et al., 2007; Oostenveld et al., 2011; Smith et al., 2004; Quinn et al., 2022). The beamforming procedure consisted of two main steps: construction of a forward model describing the projection of source activity to the MEG sensors, and computation of an inverse solution used to estimate source-level activity over time **(Figure 1d)**.

The forward model treated each brain source as an active dipole and estimated how unitary dipole activity would be reflected across the MEG sensors. Individual structural T1 images were co-registered to the MEG data using the digitised head shape and fiducial landmarks. A single-shell head model was then used to compute the leadfield. Following the original

FREQ-NESS pipeline, source reconstruction was performed using the magnetometer channels and an 8-mm whole-brain grid, yielding 3559 dipole locations throughout the brain. This choice was motivated by the sensitivity of magnetometers to deeper brain activity and by the use of a consistent sensor type for beamforming. For participants without an individual structural T1 scan, the MNI152-T1 template at 8-mm spatial resolution can be used for leadfield computation.

Leadfield models were initially computed for three principal orientations at each dipole location and then reduced to a single orientation using singular value decomposition. Beamformer weights were estimated from the sensor covariance matrix computed over the resting-state epoch and were normalised to reduce depth bias and overfitting. Applying these weights to the preprocessed MEG signal yielded one neural activity index time series for each of the 3559 brain voxels **(Figure 1e)**. These source-reconstructed voxel time series formed the input to the FREQ-NESS analysis.

### Frequency-resolved network estimation using FREQ-NESS

Frequency-resolved brain networks were estimated using FREQ-NESS (Rosso et al., 2025), a multivariate source-separation approach based on generalized eigendecomposition (GED) **(Figure 1e-h)**. FREQ-NESS identifies, at each frequency, weighted combinations of source-level voxel time series that maximises narrowband covariance relative to broadband covariance. Beyond quantifying the prominence of frequency-specific components, the method estimates their associated spatial activation patterns, thereby characterizing both the temporal and spatial organization of brain networks.

For each participant, the input to FREQ-NESS was a source-level data matrix arranged in a voxels-by-time format **(Figure 1e)**. The analysis was repeated across a dense frequency grid spanning approximately 0.2 to 97.6 Hz. To facilitate replication, the grid followed the original FREQ-NESS implementation and was anchored around 2.439 Hz, with six frequencies sampled below and 80 frequencies sampled above this reference value, resulting in 86 frequency bins (Rosso et al., 2025). In the present resting-state ageing analysis, this value was used only as a reference for constructing the frequency grid and filter widths. The filter full-width at half maximum was anchored at 0.35 Hz and scaled logarithmically (base 10) across the frequency range.

For each frequency, two covariance matrices were computed from the source-level voxel data **(Figure 1f)**. The first was a broadband reference covariance matrix, computed from the unfiltered voxel time series. The second was a narrowband covariance matrix, computed after filtering the same voxel time series around the frequency of interest using Gaussian-shaped filters in the frequency domain. The narrowband covariance matrix was stabilised by adding a small identity perturbation, and the broadband covariance matrix was regularised using shrinkage, with the shrinkage factor set to 0.01.

Generalized eigendecomposition (GED) was then used to identify spatial filters that maximised narrowband covariance relative to broadband covariance **(Figure 1g)** (Baliviera et al., 2025; Cohen, 2017, 2022; Moumdjian et al., 2025; Rosso et al., 2023, 2026). For each frequency, the resulting eigenvectors defined the spatial filters, and the corresponding eigenvalues quantified the amount of variance explained by each component. Eigenvalues were sorted in descending order and normalised by the sum of all eigenvalues, yielding the percentage of variance explained by each component. The full eigenspectrum was computed at each frequency, while the leading components were retained for subsequent spatial and statistical analyses. The first GED component was used as the main component of interest because it captured the dominant frequency-specific network at each frequency.

Together, the eigenvalue distribution across frequencies and the corresponding spatial activation patterns defined the participant-specific network landscape **(Figure 1h-i)**. A local peak in this landscape indicates that the leading component explains a relatively large proportion of covariance at that frequency compared with the broadband reference structure, and therefore reflects stronger frequency-specific network attunement.

### Spatial activation patterns

Spatial activation patterns were computed by multiplying the filters by the narrowband covariance matrix **(Figure 1h)**. These activation patterns express the contribution of each voxel to a given frequency-specific component. Because the sign of GED-derived spatial filters is arbitrary, in order to prevent that some patterns would average out at the group level, activation patterns were interpreted in terms of their magnitude. For group-level visualisation and spatial modelling, activation maps were converted to absolute values, normalised within each participant and frequency, and thresholded at the mean plus one standard deviation of the activation values. The resulting maps were written as NIfTI images in 8-mm MNI space, preserving the voxel-wise spatial organisation of the source-reconstructed data.

### Network landscape comparisons

Our primary analysis compared the dominant frequency-resolved network landscape between younger and older adults. For each participant, we extracted the normalised eigenvalue of the first GED component at each frequency. Group means and 95% confidence intervals were then computed across participants and plotted as a function of frequency **(Figure 1i)**. This provided a frequency-resolved summary of how the prominence of the leading resting-state network differed between age groups.

For statistical testing, first-component eigenvalues were compared between younger and older adults at each frequency bin using independent-samples tests. To control for multiple comparisons across frequency bins, p-values were corrected using the Benjamini-Hochberg false discovery rate (FDR) procedure (Benjamini & Hochberg, 1995). Analyses were first carried out separately in each dataset and then repeated in the combined sample. The combined analysis was used for the main inference, while the separate-dataset analyses were used to assess robustness across independent samples.

The FREQ-NESS analysis was performed across the full frequency range up to approximately 97.6 Hz, and statistical comparisons were conducted across the complete spectrum. For visualisation, figures display frequencies up to 40 Hz to improve readability and because no significant age-related differences were detected above this range. Frequency ranges were interpreted with reference to conventional bands: delta, theta, alpha, beta, and gamma. However, the analysis itself was not based on predefined canonical bands.

### Individual alpha-network peak

To quantify age-related slowing of the dominant alpha network, we estimated an individual alpha-network peak from the first-component network landscape **(Figure 1i)**. For each participant, we identified the frequency bin with the maximum explained variance within a predefined search window (7-13Hz) centred on the dominant alpha-range peak. The frequency corresponding to this maximum was taken as the participant’s alpha-network peak. Importantly, this measure differs from conventional power-based individual alpha frequency. It does not estimate the frequency at which sensor-level or source-level power is maximal. Instead, it identifies the frequency at which the leading GED component explains the greatest proportion of covariance relative to the broadband signal. It therefore reflects the frequency of maximal whole-brain network attunement. Alpha-network peak frequencies were compared between younger and older adults using Wilcoxon rank-sum tests.

### Alpha-to-beta decay of the leading network

We next quantified the decay of the leading network landscape from the individual alpha-network peak into the beta range. For each participant, first-component eigenvalues were extracted from the participant-specific alpha-network peak across the subsequent five frequency bins, thereby capturing the initial alpha-to-beta transition. To assess the robustness of this effect, the analysis was repeated using progressively longer fitting windows extending up to the subsequent 13 frequency bins. The upper limit of 13 bins was determined by the participant with the highest individual alpha-network peak, ensuring that the fitted decay did not extend beyond 30 Hz, which is the conventional upper boundary of the beta band. An exponentially decaying function was fitted to each segment of the landscape:

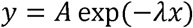

where *y* denotes explained variance, *A* is the fitted amplitude, *x* denotes frequency-bin position, and lambda (λ) is the decay coefficient. The model was fitted after linearising the function in log space:

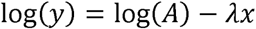

The decay coefficient λ was used as a compact index of how sharply the leading alpha-network component declined into the beta range. Larger positive values indicate a steeper decay, whereas smaller values indicate a flatter alpha-to-beta profile. Group differences in decay coefficients were assessed using Wilcoxon rank-sum tests.

### Entropy of the component landscape

To assess how concentrated or distributed the network landscape was across components, we computed quadratic Rényi entropy from the eigenvalue spectrum at each frequency. The normalised eigenvalues were treated as a probability distribution over GED components. For each participant and frequency, quadratic Rényi entropy was computed as:

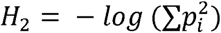

where *p_i_* is the normalised eigenvalue of component *i*. Lower entropy indicates that variance is concentrated in a smaller set of dominant components, whereas higher entropy indicates a more even distribution of variance across components. Infinite entropy values were treated as missing values.

Entropy was computed across frequencies for each participant. Group means and 95% confidence intervals were then estimated separately for younger and older adults. Younger and older adults were compared at each frequency using Wilcoxon rank-sum tests, with FDR correction applied across frequency bins.

### Spatial gradients of frequency-resolved networks

Finally, we tested whether ageing altered the spatial organisation of the leading frequency-resolved networks. For each participant, frequency, and spatial dimension, we extracted the MNI coordinates of suprathreshold (above mean + standard deviation) voxels from the first-component activation pattern. We then computed a centre of mass for each participant’s activation pattern at each frequency. This allowed us to assess whether the dominant activation pattern shifted systematically along the left-right, posterior-anterior, or inferior-superior axes as a function of frequency.

Spatial organisation was first inspected across the frequency spectrum and then summarised within canonical frequency ranges. Within each frequency range, spatial gradients were modelled separately for the x, y, and z MNI axes. A linear polynomial model was fitted to describe how the component centre of mass varied as a function of the frequency index of dominant activation. We investigated both the slope and intercept of the polynomial to characterise the gradient and spatial shifts. Group differences in spatial-gradient coefficients were then tested between younger and older adults. This analysis was designed to determine whether ageing was associated with systematic reorganisation of the spatial configuration of dominant resting-state networks, complementing the spectral analyses of alpha-network slowing, alpha-to-beta decay, and entropy.

### Statistical analysis

All statistical analyses were performed in MATLAB. The first-component eigenspectrum was compared between younger and older adults at each frequency bin, with false discovery rate correction applied across frequencies. Individual alpha-network peak, alpha-to-beta decay coefficients, entropy values, and spatial-gradient parameters were compared between groups using non-parametric Wilcoxon rank-sum tests unless otherwise specified. Frequency-wise analyses were corrected for multiple comparisons using false discovery rate correction. For visualisation, group means were plotted with 95% confidence intervals.

Analyses were conducted separately within each dataset and in the combined sample. The separate-dataset analyses were used to evaluate reproducibility across independent cohorts, whereas the combined sample was used for the primary statistical inference.

## Results

We applied FREQ-NESS to source-reconstructed resting-state MEG data from 164 healthy participants drawn from two independent datasets. FREQ-NESS uses generalized eigendecomposition to identify frequency-specific components of whole-brain covariance and their corresponding spatial activation patterns across a defined frequency grid. We then examined age-related differences using five complementary analyses: (1) network landscape comparisons, based on the normalized eigenvalue of the leading GED component at each frequency; (2) the individual alpha-network peak; (3) alpha-to-beta decay of the leading network; (4) entropy of the component landscape; and (5) spatial gradients of frequency-resolved networks.

### Ageing reshapes the dominant resting-state network landscape

The network landscape, namely the distribution of the leading eigenvalue across frequencies, showed a clear resting-state profile in both age groups **(Figure 2a)**. Across participants, the network landscape was characterized by high explained variance at low frequencies and a prominent local maximum in the alpha range. This overall structure closely resembled the resting-state profiles reported in previous FREQ-NESS studies on both younger and older participants (Rosso et al., 2025; Malvaso et al., 2025).

**Figure 2.**
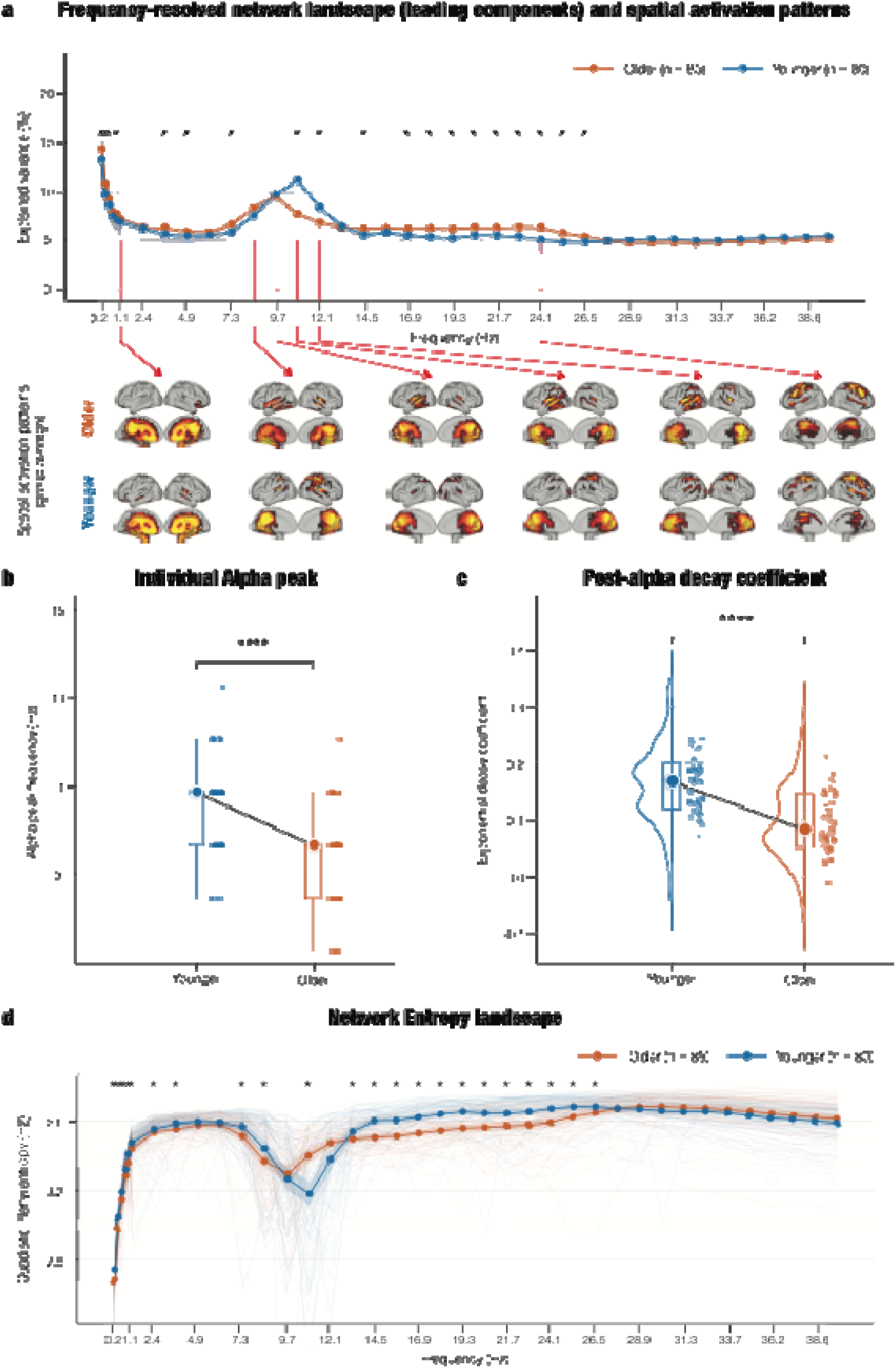
Age-related differences in the frequency-resolved network landscape. **(a)** Normalised eigenvalues of the leading GED components, expressed as percentage of explained variance, across frequencies in younger and older adults. Thin lines show individual participant landscapes, thick lines and points show group means, and shaded regions show 95% confidence intervals. The lower-frequency portion of the analysed spectrum is displayed. Asterisks indicate frequency bins at which the groups differed after FDR correction across frequencies. Compared with younger adults, older adults showed a lower-frequency and less sharply expressed alpha-range maximum, together with a less steep decay from the alpha peak into the beta range. **(b)** Group-averaged spatial activation patterns of the leading GED component at representative frequencies indicated by the red guidelines in panel a. The upper and lower rows show the corresponding activation patterns for older and younger adults, respectively. Maps represent the magnitude of the normalised, thresholded (mean + 1 standard deviation) activation patterns, with brighter yellow colours indicating a greater contribution to the frequency-specific component. **(c)** Distribution of individual alpha-network peak frequencies, defined for each participant as the frequency at which the leading GED component explained the greatest proportion of variance within the predefined alpha-range search window. **(d)** Distribution of post-alpha exponential decay coefficients of the alpha-to-beta transition, estimated per participant from the alpha-network peak into the beta range. Larger positive coefficients indicate a steeper decline in leading eigenvalues, whereas smaller coefficients indicate a flatter post-alpha distribution. In panels c and d, points represent individual participants, box-and-whisker elements summarise the group distributions, and large circles show the group means. **(e)** Quadratic Rényi entropy calculated from the complete normalised eigenspectrum at each frequency. Lower entropy indicates that covariance is concentrated in a smaller effective set of components, whereas higher entropy indicates a more even distribution across components. Thin lines show individual participants, and thick lines and shaded regions show group means and 95% confidence intervals, respectively. Asterisks indicate frequency bins with significant group differences after false discovery rate correction. Ageing shifted and broadened the alpha-range entropy minimum and produced frequency-dependent differences extending across the alpha-to-beta transition. Orange denotes older adults and blue denotes younger adults throughout.

Despite this shared overall structure, younger and older adults showed both local and global differences in the shape of the dominant network landscape. In younger adults, the alpha-range maximum was more sharply expressed and occurred at a higher frequency. In older adults, the alpha maximum was shifted toward lower frequencies (Wilcoxon rank-sum test: W = 8415, z = 4.63, *p < .001*) and the exponential decay from alpha into beta frequencies was less steep (Wilcoxon rank-sum test: W = 8868, z = 5.9, *p < .001*). This produced a flatter network landscape in older adults across the alpha-to-beta range, indicating reduced attunement of the dominant resting-state network.

Frequency-wise group comparisons, conducted across frequencies up to 90 Hz, confirmed significant age-related differences in the first-component eigenspectrum at 19 frequency bins: 0.20, 0.41, 0.61, 1.02, 3.64, 4.85, 7.26, 10.87, 12.07, 14.48, 16.89, 18.09, 19.30, 20.50, 21.70, 22.91, 24.11, 25.32, and 26.52 Hz (FDR-corrected q = 0.0057). These effects were most evident around the alpha peak and the subsequent alpha-to-beta transition. The same pattern was present when the two datasets were inspected separately, indicating that the combined-sample result was not driven by a single cohort.

Together, these findings show that ageing alters the overall shape of the dominant resting-state network landscape. The effect was not limited to a simple reduction in explained variance at one frequency but involved a broader reorganisation of brain networks as a function of frequency.

### The dominant alpha-network peak slows with age

We next tested whether the shift in the alpha-range maximum reflected a reliable slowing of the dominant alpha network. For each participant, we identified the frequency at which the first GED component explained most variance within the alpha range. This provided an individual alpha-network peak, reflecting the frequency of maximal network attunement rather than a conventional power-based individual alpha frequency.

Older adults showed a clear shift toward lower alpha-network peak frequencies compared with younger adults **(Figure 2c)**. The distribution of individual alpha-network peaks were slowed in the older group, indicating that the dominant resting-state network reached its maximum explained variance at a lower frequency in ageing. This group difference was significant (Wilcoxon rank-sum test: W = 8415, z = 4.63, *p < .001*)

This result provides a network-level counterpart to the well-established slowing of individual alpha frequency in healthy ageing. Importantly, the effect was observed in the frequency-resolved covariance structure of the source-reconstructed whole-brain signal, indicating that alpha slowing was expressed at the level of dominant resting-state network organisation rather than only in local or sensor-level power.

### Alpha-to-beta decay flattens with age

The network landscape suggested that ageing affected not only the location of the alpha peak in the spectrum, but also the way the dominant alpha network declined into the beta range. To quantify this effect, we fitted an exponential decay function to the leading eigenvalue distribution, beginning at each participant’s alpha-network peak and extending into the alpha-to-beta transition. The resulting decay coefficient provided a compact measure of how sharply the leading eigenvalues decreased after the alpha maximum.

Older adults showed significantly smaller decay coefficients than younger adults (**Figure 2d**; W = 8868, z = 5.9, *p < .001*). In other words, the dominant network landscape decayed less steeply from alpha into beta frequencies in older age. This flattening was visible at the group level and was also reflected in the distribution of individual decay coefficients.

This result complements the alpha-peak analysis. Whereas the individual alpha-network peak captured the shift of the dominant maximum toward lower frequencies, the decay analysis showed that ageing also changed the shape of the surrounding network landscape. The older group did not simply show a displaced alpha peak; rather, alpha activity was less sharply tuned around its peak, resulting in a less steep transition into beta frequencies and indicating that comparatively broader-band activity dominated this region of the spectrum. This pattern is consistent with reduced spectral attunement of the dominant resting-state network in ageing.

### Ageing alters the entropy of the network landscape

We then asked whether ageing changed how variance was distributed across the broader set of GED components within frequencies. To this end, we computed quadratic Rényi entropy from the normalized eigenvalue spectrum at each frequency. Lower entropy indicates that variance is concentrated in a smaller number of dominant components, whereas higher entropy indicates that explained variance is more evenly distributed across components.

Both younger and older adults showed a structured entropy landscape across the frequency spectrum **(Figure 2e)**. Entropy was not constant across frequencies, indicating that the dimensional structure of the component landscape varied systematically as a function of frequency. In particular, the alpha range showed a marked change in entropy, consistent with the strong dominance of a reduced set of networks around the alpha peak.

Ageing altered this entropy landscape in a frequency-dependent manner. Group differences were observed across frequencies up to 90 Hz, with significant effects at 0.20, 0.41, 0.61, 1.02, 7.26, 10.87, 13.28, 14.48, 15.68, 16.89, 18.09, 19.30, 20.50, 21.70, 22.91, 24.11, and 25.32 Hz after correction for multiple comparisons (FDR-corrected critical p = 0.0016). The strongest differences were centred around the alpha range and the alpha-to-beta transition. In older adults, the entropy profile was less sharply organised around the alpha peak and showed a more gradual transition across neighbouring frequencies.

These results indicate that ageing affects not only the dominant component itself, but also the distribution of variance across the wider component landscape. Rather than reflecting a uniform increase or decrease in entropy across the spectrum, the effect was frequency-specific. This suggests that ageing changes how resting-state covariance is organized into dominant and secondary network components, particularly around frequencies where the leading network is normally most strongly expressed.

### Spatial topology of frequency-resolved networks changes with age

Finally, we tested whether spectral changes in the network landscape were accompanied by changes in spatial topology **(Figure 2b)**. For each participant and frequency, we extracted spatial summaries of the leading component’s activation pattern in MNI space and modelled age-related differences along the left-right, posterior-anterior, and inferior-superior axes. These analyses were performed across the frequency spectrum and then summarized within canonical frequency ranges.

Ageing was associated with band-specific shifts in the spatial configuration of the dominant networks **(Figure 3a and 3b)**. Frequency-wise comparisons across frequencies up to 42 Hz revealed a significant group difference along the left-right axis at 8.46 Hz (FDR-corrected critical p = 0.0120). Along the posterior-anterior axis, significant differences were observed at 1.02, 3.64, 4.85, 6.05, 12.07, 14.48, 15.68, 16.89, 18.09, 19.30, 21.70, 25.32, and 26.52 Hz after FDR correction. Along the inferior-superior axis, significant differences were observed at 0.81, 6.05, 7.26, 8.46, 9.66, 10.87, 14.48, 15.68, 16.89, 18.09, 24.11, 25.32, and 26.52 Hz after FDR correction. These spatial effects were not uniform across the frequency spectrum. In the theta range, younger adults showed more anterior and superior spatial estimates than older adults. In the alpha range, younger adults showed more rightward, posterior, and superior spatial estimates, indicating that the dominant alpha network in older adults was shifted away from the posterior-superior configuration observed in younger adults. In the beta range, younger adults showed more posterior and inferior spatial estimates, whereas older adults showed a relative shift toward more anterior and superior beta-band activation patterns.

**Figure 3.**
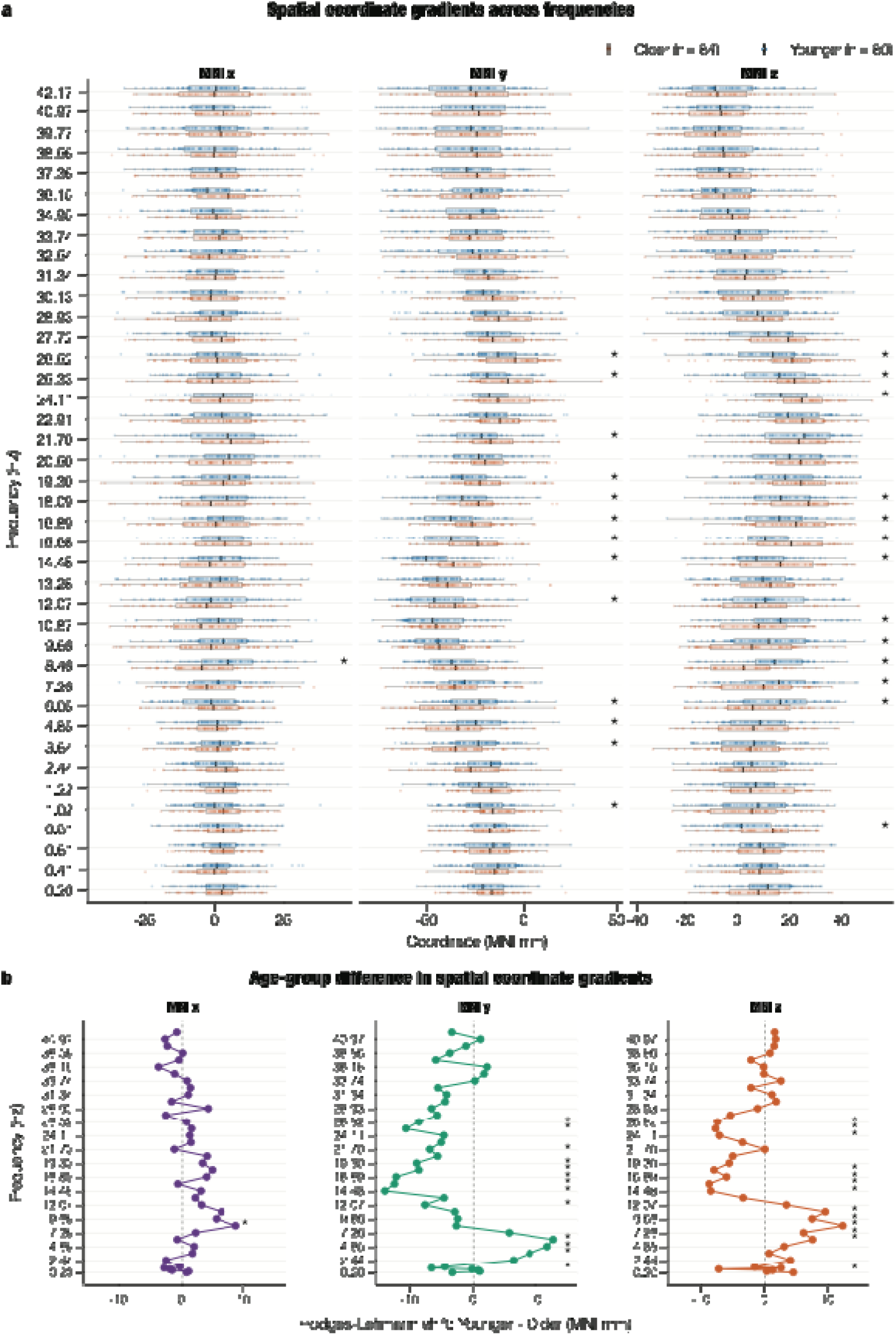
Age-related differences in the spatial coordinate gradients of the leading frequency-resolved network. **(a)** Spatial centre-of-mass coordinates of spatial patterns across frequencies from 0.20 to 42.17 Hz. For each participant and frequency, the centre of mass was calculated from the suprathreshold (above mean and standard deviation) voxels of the corresponding first-component activation pattern. Distributions are shown separately for the MNI (x), (y), and (z) axes, representing the left-right, posterior-anterior, and inferior-superior dimensions, respectively. Positive (x) coordinates indicate a rightward position, positive (y) coordinates indicate an anterior position, and positive (z) coordinates indicate a superior position. Points represent individual participants, and box-and-whisker elements summarise the group distributions at each frequency. Orange denotes older adults and blue denotes younger adults. Asterisks indicate frequencies at which younger and older adults differed significantly after false discovery rate correction across frequencies. **(b)** Hodges-Lehmann estimates of the age-group shift in centre-of-mass coordinates at each frequency, calculated as younger minus older adults. Positive values indicate that the leading activation pattern was positioned more rightward, anteriorly, or superiorly in younger adults for the (x), (y), and (z) axes, respectively; negative values indicate the corresponding relative shift in older adults. The vertical dashed line denotes no group difference. Asterisks identify frequencies showing significant group differences after false discovery rate correction.

The spatial pattern therefore suggests that ageing reshapes the anatomical configuration of frequency-resolved resting-state networks. The effect was strongest in the same broad frequency range where the spectral landscape showed alpha slowing and alpha-to-beta flattening. This convergence indicates that spectral and spatial changes were not independent observations, but complementary aspects of a broader reorganisation of resting-state network structure.

## Discussion

Using a minimally constrained, data-driven framework, we provide an integrated account of how healthy ageing reshapes frequency-resolved whole-brain organisation. Within a single analysis, we showed how ageing altered the prominence and spectral profile of dominant networks, the entropy across networks, and their spatial expression, with effects varying systematically across frequencies. These findings reveal a selective reorganisation of the resting-state network landscape rather than a uniform loss of network integrity, and position classical alpha slowing as one manifestation of a broader change in whole-brain functional organisation.

The slower alpha-network peak provides the clearest connection with established electrophysiological correlates of ageing. A reduction in individual alpha frequency and power is among the most reproducible resting-state EEG and MEG findings in healthy older adults, although its estimated magnitude depends on age range, recording conditions, and peak-detection procedures (Cassani et al., 2019; Fernández-Rubio et al., 2025; Park et al., 2024; Ishii et al., 2017; Sahoo et al., 2020; Scally et al., 2018; Merkin et al., 2023). Importantly, FREQ-NESS moves beyond detecting changes in peaks in the power spectrum, allowing us to quantify the slowing in alpha activity at the level of whole-brain networks.

Beyond the shift in peak frequency, older adults showed reduced spectral concentration of network prominence around the alpha maximum, expressed as a flattened decay into the beta range. Recent lifespan MEG studies have reported age-related changes in beta power, bursts, and connectivity alongside weakening or slowing of alpha activity, with the direction of beta effects depending on cortical system, age range, and measure (Gohil et al., 2026; Ranasinghe et al., 2025; Ruuskanen et al., 2026; Sahoo et al., 2020). Power-based analyses across the frequency spectrum similarly suggest that ageing alters the relative expression of alpha- and beta-range activity rather than producing a uniform spectral reduction (Quinn et al., 2025). Although these measures are not directly equivalent, they support an age-related rebalancing of alpha- and beta-range organisation, for which our decay coefficient provides a network-level index.

In order to extend the current analysis beyond the leading GED component, we analysed quadratic Rényi entropy across normalised eigenvalues at each frequency. Lower values indicate that variance is explained by a smaller set of effective components, whereas higher values indicate a more even distribution across a larger component set. The direction of the ageing effect differed between the alpha and beta ranges. This pattern argues against a global increase or decrease in entropy with age and instead indicates a frequency-dependent redistribution of the variance explained across components. This is consistent with previous work showing that age-related changes in neural complexity and variability depend on the scale at which they are measured. For example, multiscale entropy analyses have demonstrated timescale-dependent alterations in brain signal variability with ageing (McIntosh et al., 2014; Ando et al., 2022), while other entropy and complexity measures have identified frequency-specific or region-specific changes in electrophysiological dynamics (Alù et al., 2021; Shumbayawonda et al., 2020). Related multivariate measures, including effective rank, participation ratio, and principal-component entropy, likewise describe how covariance is distributed across concurrent components (Diambra et al., 2026; Roy & Vetterli, 2007). The frequency dependence observed here may help reconcile heterogeneous findings from temporal entropy studies of ageing, which vary across estimators, scales, and frequency ranges (Cacciotti et al., 2024). Healthy ageing may weaken the sharply alpha-centred organisation of whole-brain covariance and redistribute network concentration across a broader alpha-to-beta regime.

Spatial reconfiguration was similarly dependent on frequency and spatial axis, with no evidence for a uniform posterior-to-anterior or otherwise unidirectional shift in older adults. This is consistent with source-level electrophysiological evidence that posterior alpha generators and sensorimotor beta systems follow partly distinct ageing trajectories (Park et al., 2024; Perinelli et al., 2022), and can be considered in relation to the cortical hierarchy of intrinsic frequencies (Mahjoory et al., 2020). Although the posterior-anterior shift in ageing provides a useful conceptual comparison, this framework was developed primarily from task-evoked fMRI (Davis et al., 2008). Resting-state centre-of-mass differences therefore do not establish compensatory recruitment, which would require evidence that altered activity responds to neural demand and supports preserved behaviour (Cabeza et al., 2018). The observed centroids may instead reflect altered weighting of posterior sensory, anterior association and sensorimotor generators, as well as changes in the spatial extent of activation, or structural and physiological constraints on source expression.

Taken together, the network landscape, network entropy, and spatial reorganisation of network activation patterns provide complementary descriptions of frequency-resolved whole-brain organisation. Across these three domains, ageing was associated not with a uniform loss of network organisation, but with a selective and frequency-dependent reconfiguration of spectral prominence, component concentration, and spatial expression. Reduced functional segregation and modularity, greater between-system coupling, altered efficiency and changes in structural-functional correspondence have been reported across resting-state studies (Chan et al., 2014; Deery et al., 2023; Jauny et al., 2024), but these graph-based measures are not equivalent to GED eigenspectrum concentration. Likewise, changes in dynamic-state occupancy or repertoire describe temporal switching between configurations rather than the static covariance distribution estimated here. Electrophysiological evidence is also not uniform: Coquelet et al. (2017), for example, reported largely preserved static and dynamic envelope-connectivity organisation in carefully screened healthy older adults. Such null findings argue against a universal loss of network integrity and support a more selective interpretation. Together, current findings argue against a universal decline in network integrity and instead support the view that healthy ageing selectively reshapes how whole-brain activity is organised across frequencies and space. Because conventional network estimates are sensitive to choices concerning frequency bands, parcellation, connectivity metrics, and preprocessing, the present data-driven framework provides a complementary and more integrated description of this reorganisation.

Several biological processes could potentially contribute to this reorganisation. Ageing affects axonal and myelin integrity, synaptic regulation, neuromodulatory systems, and metabolic support, all of which can constrain the timing and coordination of large neural populations (Bishop et al., 2010; Turrini et al., 2023). Changes in conduction delay provide one possible link between slower network rhythms and altered large-scale coordination. In a lifespan MEG and computational-modelling study, Pathak et al. (2022) proposed that increased coupling could partially preserve alpha-range phase synchrony despite longer transmission delays and slower oscillatory frequency. Altered thalamocortical timing, synaptic gain, excitatory-inhibitory regulation and neuromodulatory tone could also influence both preferred frequency and the relative dominance of concurrent components (Steriade et al., 1990; McCormick & Bal, 1997; Voytek et al., 2015; Werkle-Bergner et al., 2022). Determining how these cellular and circuit-level mechanisms give rise to the frequency-resolved network reorganisation observed here will require future multimodal studies combining electrophysiology with measures of structural, molecular and neurochemical ageing.

Methodologically, FREQ-NESS provides a complementary perspective relative to conventional spectral and network analyses. It avoids defining the primary components through canonical frequency bands or anatomical regions and identifies simultaneous whole-brain covariance patterns that are prominent at specific frequencies relative to broadband structure (Rosso et al., 2025). This differs from power analysis, periodic-aperiodic parameterisation, pairwise connectivity, graph construction, and temporal entropy. While each of these approaches addresses a distinct aspect of neural organisation and remains useful to discuss our findings, this work complements large source-MEG studies based on canonical power analyses, coherence and dynamic states (Gohil et al., 2026), frequency-resolved connectivity (Ruuskanen et al., 2026), later-life periodic and aperiodic activity (Ranasinghe et al., 2025), and cross-dataset spectral replication (Quinn et al., 2025). The analysis of two independent datasets further strengthens confidence that the principal pattern was not unique to one cohort.

By providing an integrated account of spectral and spatial reorganisation in healthy ageing, the present study establishes a foundation for several important directions of future research. First, the cross-sectional contrast between categorical younger and older groups cannot reveal within-person change, distinguish linear from nonlinear lifespan trajectories, or determine whether the effects precede, accompany, or follow cognitive change. Continuous-age and longitudinal designs are required to establish ageing trajectories. Second, despite the strongest prominence of the leading component, other components might also give relevant functional insight. While in the current study we capitalised on the entire set of components to compute entropy measures, future work should further investigate relationships between components and the higher-order organisation of brain dynamics, including properties such as irreversibility and non-equilibrium interactions (Nartallo-Kaluarachchi et al., 2025).

In conclusion, healthy ageing altered more than a dominant alpha rhythm. The slowing down of the alpha-network maximum was accompanied by a flatter alpha-to-beta profile, a frequency-dependent redistribution of variance explained across GED components, and reconfiguration of the spatial patterns of each network. Alpha slowing can therefore be situated within a wider change in the frequency-resolved architecture of endogenous whole-brain dynamics. While the framework presented in this study does not establish deterioration, compensation or clinical relevance, it provides a structured way to examine how ageing jointly affects spectral preference, component concentration, and neuroanatomy. Longitudinal, behaviourally informed, and multimodal studies are needed to determine whether these network-landscape features track individual ageing trajectories, cognitive maintenance or vulnerability to pathological ageing.

## Code availability

The codes are available at the following link: https://github.com/cognikita-cn/FreqNESS_HealthyAgeing

The FREQNESS Toolbox for Matlab and Python is available in the following Github repository: https://github.com/mattiaRosso92/Frequency-resolved_brain_network_estimation_via_source_separation_FREQ-NESS/tree/main/FREQNESS_Toolbox

## Data availability

The neuroimaging data related to the experiment is available upon reasonable request.

## Acknowledgements

The Center for Music in the Brain (MIB) is funded by the Danish National Research Foundation (project number DNRF117), The Lundbeck Foundation (R469-2024-1573) and Købmand Herman Sallings Fond.

L.B. is supported by Sapere Aude: Independent Research Fund Denmark (DFF) Research Leader (grant ID: 10.46540/5253-00003B), Lundbeck Foundation (Talent Prize 2022), Carlsberg Foundation (CF20-0239), Center for Music in the Brain, Linacre College of the University of Oxford and Nordic Mensa Fund.

Mattia Rosso is supported by Center for Music in the Brain and Nordic Mensa Fund

MLK is supported by Center for Music in the Brain and Centre for Eudaimonia and Human Flourishing, which is funded by the Pettit and Carlsberg Foundations.

We thank Emma Risgaard Olsen, Mathias Klarlund, Orla Mallon, Francesco Carlomagno, Luna Frausing and Victor Pando-Naude for their important contribution during the data collection.

## Author contributions

L.B, M.R. and N.K. conceived the initial hypotheses and methodological development. Resources were recruited by L.B., M.L.K., E.S. and P.V. for performing data collection and analysis. G.F.R., E.S., M.H.A. and L.B. collected the data. L.B. and G.F.R. performed pre-processing. N.K., M.R. and L.B. developed the analytical pipeline. M.L.K., P.V., M.H.A., G.F.R., L.B. and M.R. provided essential help to interpret and frame the results within the neuroscientific and analytical literature. N.K. wrote the first draft of the manuscript, which was primarily integrated by L.B. and M.R. The figures were prepared by N.K., with the support of L.B. and G.F.R. All the authors contributed to and approved the final version of the manuscript.

